# Automated Cytogenetic Biodosimetry at Population-Scale

**DOI:** 10.1101/718973

**Authors:** PK Rogan, R Lu, E Mucaki, S Ali, B Shirley, Y Li, R Wilkins, F Norton, O Sevriukova, D Pham, E Ainsbury, J Moquat, R Cooke, T Peerlaproulx, E Waller, JHM Knoll

**Author notes:** Correspondence: Peter K. Rogan, University of Western Ontario London, Ontario N6A 5C1 Canada, E. CrediT [Contributor Roles Taxonomy; www.casrai.org]: Conceptualization (Rogan), Data curation (Rogan, Ali, Lu, Shirley, Li), Formal analysis (Rogan, Mucaki, Lu, Ali, Shirley), Funding (Rogan, Waller, Knoll), Investigation (Rogan, Lu, Ali, Cooke, Peerlaproulx), Methodology (Rogan, Lu, Mucaki, Ali, Shirley), Project administration (Rogan, Waller, Knoll), Resources (Wilkins, Norton, Sevriukova, Pham, Ainsbury, Moquat, Cooke, Peerlaproulx), Software (Lu, Ali, Shirley, Li), Supervision (Rogan, Waller, Knoll), Validation (Lu, Mucaki), Visualization (Lu, Mucaki, Shirley), Writing (Rogan, Lu, Mucaki, Knoll).

## Abstract

**Introduction:** The dicentric chromosome (DC) assay accurately quantifies exposure to radiation, however manual and semi-automated assignment of DCs has limited its use for a potential large-scale radiation incident. The Automated Dicentric Chromosome Identifier and Dose Estimator Chromosome (ADCI) software automates unattended DC detection and determines radiation exposures, fulfilling IAEA criteria for triage biodosimetry. We present high performance ADCI (ADCI-HT), with the requisite throughput to stratify exposures of populations in large scale radiation events.

**Methods:** ADCI-HT streamlines dose estimation by optimal scheduling of DC detection, given that the numbers of samples and metaphase cell images in each sample vary. A supercomputer analyzes these data in parallel, with each processor handling a single image at a time. Processor resources are managed hierarchically to maximize a constant stream of sample and image analysis. Metaphase data from populations of individuals with clinically relevant radiation exposures after simulated large nuclear incidents were analyzed. Sample counts were derived from US Census data. Analysis times and exposures were quantified for 15 different scenarios.

**Results:** Processing of metaphase images from 1,744 samples (500 images each) used 16,384 CPUs and was completed in 1hr 11min 23sec, with radiation dose of all samples determined in 32 sec with 1,024 CPUs. Processing of 40,000 samples with varying numbers of metaphase cells, 10 different exposures from 5 different biodosimetry labs met IAEA accuracy criteria (dose estimate differences were < 0.5 Gy; median = 0.07) and was completed in ~25 hours. Population-scale metaphase image datasets within radiation contours of nuclear incidents were defined by exposure levels (either >1 Gy or >2 Gy). The time needed to analyze samples of all individuals receiving exposures from a high yield airborne nuclear device ranged from 0.6-7.4 days, depending on the population density.

**Conclusion:** ADCI-HT delivers timely and accurate dose estimates in a simulated population-scale radiation incident.

## Introduction

Radiation emergency management in a nuclear radiation incident over a large geographic region or affects many individuals will involve an extraordinary degree of coordination between first responders, testing laboratories and clinical personnel. The scope of testing in population-based scenarios has been recognized to exceed the capacity most biodosimetry laboratories. Not only would the volumes of samples overwhelm these laboratories, but the impact of large volume testing on accuracy has not been established.

The dicentric chromosome assay (DCA, [1]) is the gold standard test within the clinically relevant and treatable radiation exposure range. While rapid tests are under development for triage purposes [2–7], the calibration curves for these assays tend to exhibit high variance, which impacts the confidence of the estimated dose. This can possibly lead to overtesting of worried well or inadequate testing of at risk, exposed populations.

Several international exercises and proposals have envisioned cooperative biodosimetry testing could be distributed over multiple laboratories to overcome the bottleneck in generating dose estimates for exposed populations [8–12]. There are a number of significant challenges inherent in implementing such a strategy. The degree of interlaboratory coordination required - including sample transport - may not be adequate to handle the workloads in an actual population-scale event. Furthermore, reliable internet resources connecting laboratories may not be available to collate results across international borders and interpretation would have to be standardized between laboratories.

Performance of the conventional DCA on all or most suspected cases of Acute Radiation Syndrome in a mass casualty would cause bottlenecks in sample processing, specifically cell culture, capture of metaphase cell images, and interpretation of those images. Without automation, the throughput of this test will probably not be adequate to triage patients for life saving cytokine therapies [8]. Semi-automated testing is efficient for small sample volumes, but processing and analysis of thousands of samples will surpass recommended therapeutic windows in a mass casualty [13]. Full automation of sample preparation, metaphase cell imaging, and interpretations of the DCA could substantially contribute to meeting testing capacity requirements to ensure timely administration of therapies. The Automated Dicentric Chromosome Identifier and Dose Estimator (ADCI) is a medical image processing system that leverages machine learning to analyze metaphase cell samples from different individuals, in the form of images, to identify dicentric chromosomes as an indicator of the patient’s level of radiation exposure that would then be used to determine the treatment needed, if any [14–16]. The current Windows-based implementation of the system substantially reduces the time required for laboratories to estimate radiation exposures relative to the manual and semiautomated analysis. However, in a large-scale radiation accident or mass casualty, data from many different individuals would need to be processed quickly. A bank of personal computers running ADCI may not be able to meet the throughput required to triage entire populations on the scale of a moderate sized city.

The present study investigates automation of analysis of cytogenetic data required for the DCA and radiation exposure assessment. ADCI was parallelized on a multiprocessor supercomputer to determine if performance is sufficient to handle the demands of population-scale exposures. This has the effect of simultaneously processing metaphase images from multiple samples, with the goal of expediting the analysis of the full set of samples. The advantage of parallelization is that it would provide dose of exposure information for clinical decision making for many exposed individuals at the same time. It is assumed that testing laboratories have the capacity to automate culturing of multiple blood samples, harvest and prepare slides of metaphase cells, and possess microscope systems for capture of metaphase cell images. We previously demonstrated a proof of concept implementation with a subset of the components of the ADCI system [17], indicating that the software could be adapted for large scale processing of many samples by master-slave rank-based scheduling of available computing resources across all samples and images. This was the starting point for the current implementation of the fully functional ADCI application on the IBM BG/Q supercomputer. This high-throughput ADCI (ADCI-HT) software was designed to rapidly analyze thousands of images in parallel, obtaining near identical dose predictions quickly compared to the time required via a commercial Windows desktop. Nuclear incident results were simulated for 15 US cities by calibrating radiation plumes predicted from physical dosimetry with population data. ADCI-HT was then used to analyze individuals exposed to >1Gy and >2Gy of radiation. Finally, analysis of actual international exercise data was analyzed, confirming ADCI-HT software predictions meets IAEA triage analysis requirements.

## Methods

### Underlying principles for accelerating ADCI by High-Throughput supercomputing

#### Porting ADCI for high throughput biodosimetry

The design of ADCI-HT emphasized the throughput of the software to handle many samples in a single run. The high-throughput (HT) version leverages previous analyses using the Windows version of the program that would be used to generate a calibration curve and derive optimal parameters for image model selection. The accelerate the software, the HT version lacks a graphical user interface, eliminating direct user interaction. The elements of ADCI C^++^ software were either replicated, modified or replaced in order to manage parallel analyses of samples with thousands of computer processors. Some compilation differences between Intel and PowerPC architectures used in the IBM-BG/Q supercomputer had to be addressed during implementation. The emphasis on large scale processing capability on BG/Q limits available input/output (I/O) resources and the system is not interactive. BG/Q reads and decodes compressed archives of multiple samples (typically 50-400), each consisting of at least 500 microscope cell images (TIF or PNG format) into memory. The elements of the system used by both versions are described in Figure 1, which is updated from our previously described Desktop Version of ADCI software [18]. The HT implementation replicates the 3 layers below the “Application” level of the ADCI-Desktop version (IV). Additionally, new function named “Sample::calculateDose()” was added to ADCI-HT. This function uses the previously ported classes to read the calibration curves previously generated with Desktop ADCI and the image selection models, filters the images in each sample according to these models, and estimates the radiation exposure doses.

**Figure 1.**
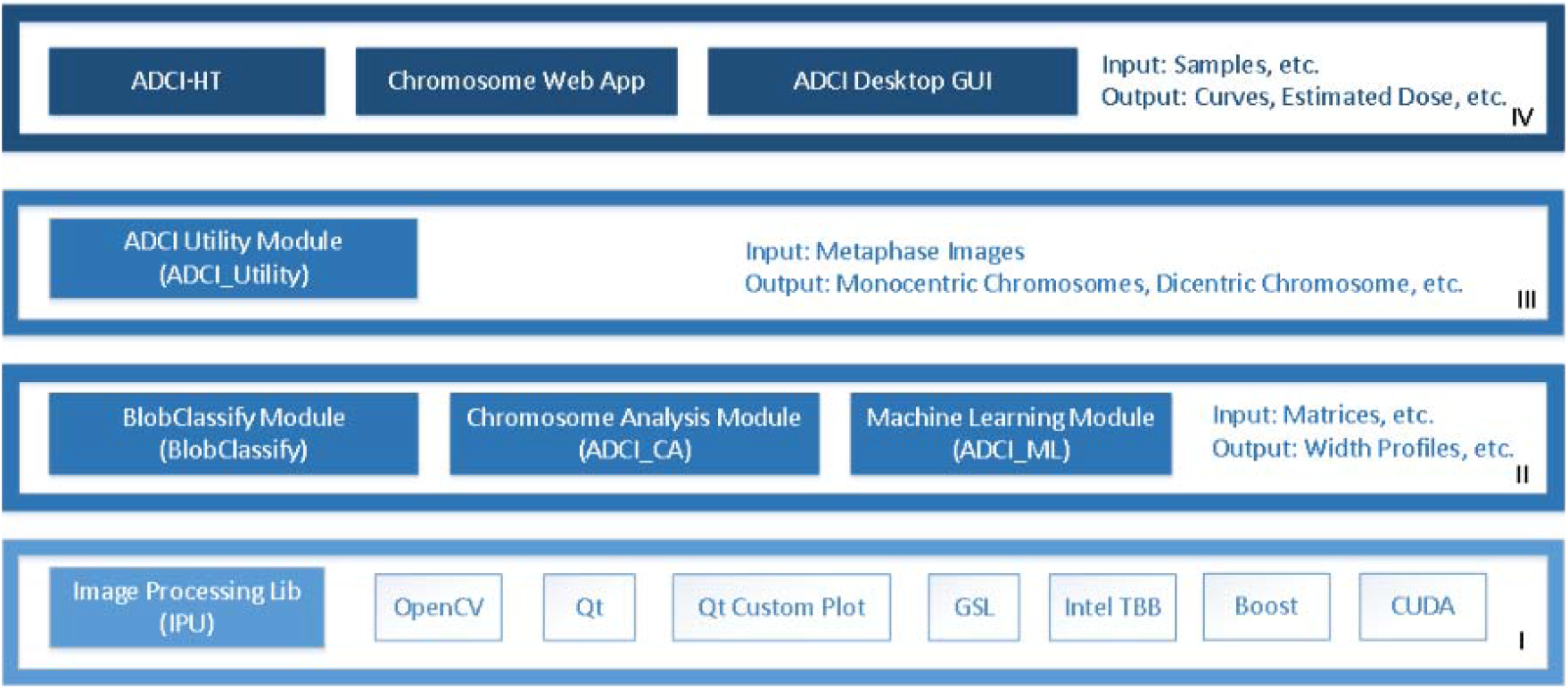
ADCI Project: Inventory of modules by software layer. Illustration of the HT implementation of the ADCI software. I. Supporting libraries: public, open source (unfilled boxes) and proprietary (filled boxes) components called by functional modules. II. Functional modules used by the ADCI Interface. III. Interface called by applications. IV. Different software Applications using ADCI interface to process images.

### Scheduling system

Efficient automated cytogenetic analysis of thousands of biodosimetry samples requires many computational tasks to be performed simultaneously. These tasks need to be scheduled to maximize concurrent use of all available processor resources to analyze all metaphase images in each sample [15]. The algorithm implemented was non-preemptive, since we assumed equivalent priority for processing all samples, and was dynamic, since the resources required depends on the number of samples and size of each sample to be processed (which cannot be known in advance).

A quantitative analysis of image processing performed by ADCI-HT used test samples from Health Canada (n=6; 540-1,136 images each) and Canadian Nuclear Laboratories (n=7; 500-1,527 images each). Performance of the scheduler was correlated with sample size (numbers of images analyzed). The time consumption to load and read a sample was correlated with the size of the compressed archive file (r=0.95) and to a lesser extent, with the number of images in the sample (r=0.65).

A data flow diagram apportions the time required to perform the tasks that ADCI performs to process metaphase images (Supplementary Figure 1). The segmentation task first takes an input image and identifies the regions containing connected objects based on pixel intensities. The chromosome analysis (CA) task identifies valid chromosomes through machine learning (ML)-based recognition of the centromere candidates and other features in each object. The identified chromosomes are then analyzed by another ML task to distinguish the dicentric chromosomes (DCs) from monocentric chromosomes (MCs). Results are then processed by the filtering task to remove false positive DCs. Finally, overall statistics are calculated for each image before the results are saved. The chromosome analysis task was found to be a primary bottleneck among all of the tasks, with the highest standard deviation for the time to process images, meaning that the requirements for this task were the most variable among all tasks. This results in CPU load imbalance, whereby some images require substantially more computing resources than others to analyze. In fact, the number of objects in each image was only weakly correlated with chromosome analysis time (r=0.07). One caveat, however, was that certain datasets created using replicate images with nearly identical sample sizes, could impact the independence, and thereby the accuracy of the statistics drawn from those samples.

Within the chromosome analysis task, individual chromosomes are segmented by local thresholding and Gradient Vector Flow (GVF; [19]) active contours. Upon extraction based on the GVF boundaries, the contour of the chromosome is partitioned using a polygonal shape simplification algorithm known as Discrete Curve Evolution (DCE) which iteratively deletes vertices based on their importance to the overall shape of the object. A Support Vector Machine (SVM) classifier selects the best set of points to isolate the telomeric regions i.e. at the ends of each chromosome. The segmented telomere regions are then tested for evidence of sister chromatid separation using a second trained SVM classifier designed to capture shape characteristics of the telomere regions and then corrected for that artifact. Afterwards, the chromosome is split into two partitions along the axis of symmetry and a modified Laplacian-based thickness measurement algorithm (called Intensity Integrated Laplacian or IIL [20]) is used to calculate the width profile of the chromosome. This profile is then used to identify a possible set of candidates for centromere location(s) and features are calculated for each of those locations. Next, another classifier is trained on expert-classified chromosomes to detect centromere locations in chromosomes. In most instances, each chromosome will contain at least one centromere. The correct centromere is generally present among the candidates. The distance from the separating hyperplane is used as an indicator for the goodness of fit of a given candidate and thereafter used to select the best candidate from the pool of candidates. Finally, we apply a machine learning method that uses image features to distinguish mono-from dicentric chromosomes. The most time-consuming sub-tasks (GVF and IIL) depend on the length of the chromosomes (correlation is 0.89), which is indicated by the area of the region of interest for each chromosome. The maximum value for GVF was high, but the average time was low, suggesting that the high value is an outlier and was not representative of all images. IIL was the actual processing bottleneck, since it required a minimum of 3 sec each to perform this step for each of the images analyzed. Dose accuracy relies on metaphase image selection criteria, which depends on thresholding and in some instances sorting images by quality and are optimized according to data generated by each biodosimetry laboratory [15].

To handle the simultaneous analysis of many samples, we developed a three-layer scheduling architecture consisting of a general manager, which directs several managers, each of which are dynamically assigned by the program to multiple worker processors (Figure 2A). The scheduler minimizes compute load imbalances between different processors. Each manager sends asynchronous requests for analysis of individual images to each worker (based on the value of a variable defined as “extraload”). Once the worker completes the analysis and submits its results, it queries managers for any outstanding requests. If a request has not been fulfilled, the worker receives that message, determines if the value of “extraload” is non-zero, then requests another image from the manager. The manager responds by sending an image while reducing the value of “extraload”. Without “extraload”, this communication would not take place at all. It is conceivable that multiple workers may receive the message before the value of the “extraload” variable is updated; however, the resources used by the resulting unnecessary processes are negligible.

**Figure 2.**
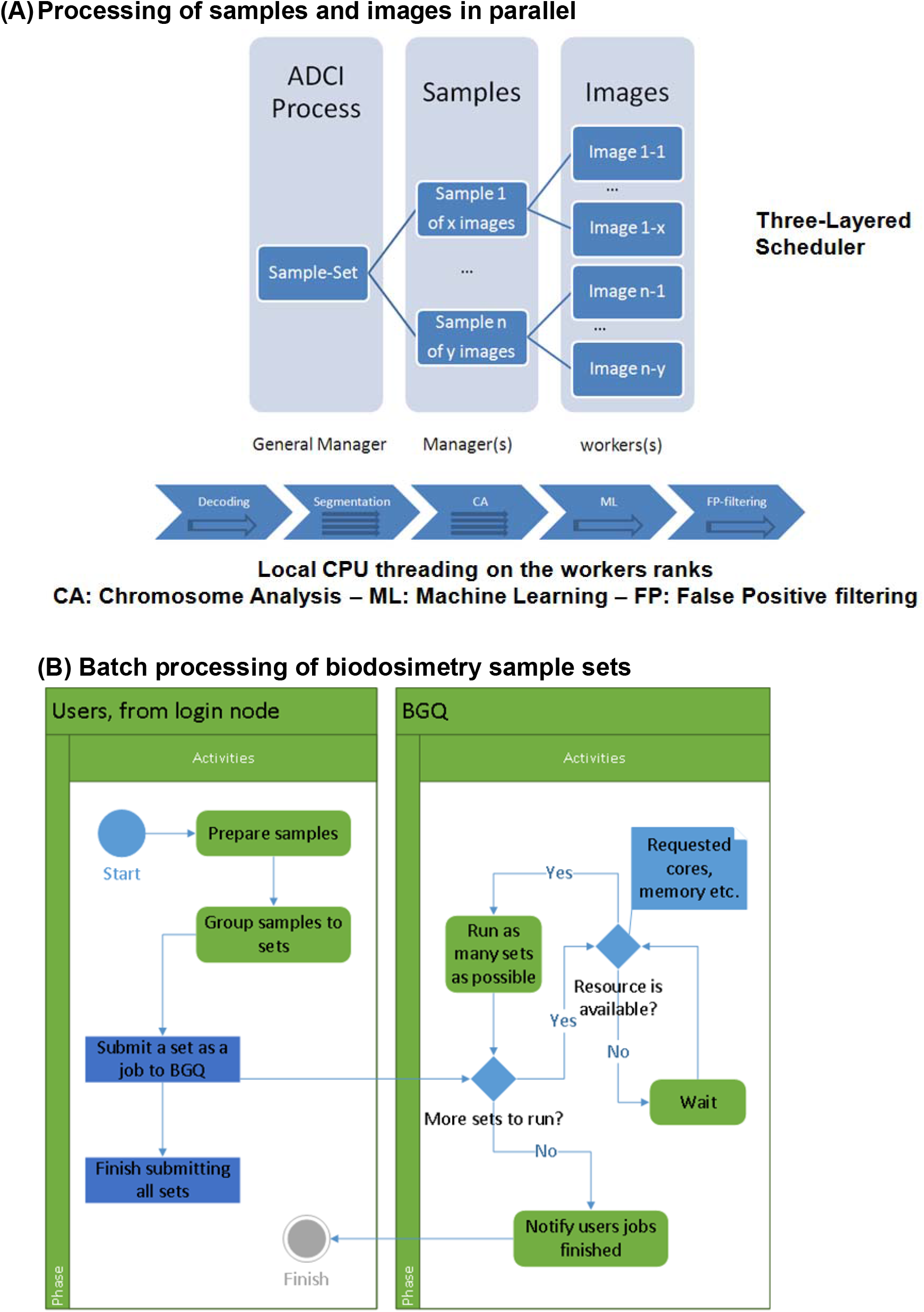
Master scheduling software in BG/Q. A) Architecture of three layer scheduler which consists of a general manager which controls individual managers which in turn are assigned multiple worker processors, minimizing compute load imbalances. B) In BG/Q, samples are grouped into sets and run on requested cores, running as many processes as possible while considering the resources available until all sets are completed.

### User interface and outputs

The ADCI-HT batch queue allocates shared compute resources to ADCI once they are available. Tasks and CPU resource requests submitted to this queue use a command line interface which provides similar functionality to the Graphical User Interface of the MS-Windows version. This interface specifies a configuration file that can assign different image selection models and calibration curves to process particular sets of samples, for example, originating from different biodosimetry laboratories. Upon completion, ADCI processed sample files are written in the same directory where unprocessed samples are located and have the same file name as the sample, appended with the suffix “.adcisample.” Dose Estimate reports are formatted as CSV files located in the directory of the samples mentioned in it. The report provides the sample name, the curve file, the selection model, the sigma value, the DC frequency and the estimated dose.

### Estimating affected population size in nuclear incidents

We estimated the size of the population exposed to different levels of radiation by intersecting geographic contours of minimum radiation exposure levels computed by HPAC v4.02 software (Hazard Prediction and Assessment Capability; developed by the Defense Threat Reduction Agency [DTRA]) with the census in these regions. KML (<u>Keyhole Markup Language</u>) based boundary location files for counties and subdivisions from the US census bureau website were downloaded in XML format. Census boundaries that overlap radiation contour plumes were obtained by transforming the polygons of each boundary to Google Maps-encoded polygons and creating javascript (‘plume-census.js’) that draws all the polygons in a map in HTML. The US Census API (<u>api.census.gov/data/2016/pep/population</u>; Application Program Interface) was interrogated with the intersecting subdivisions to estimate populations residing in the plume and the sum of all populations within a contour estimates the affected population exceeding a particular exposure level (e.g. >3, >2, >1 Gy). Applying this strategy to several large US cities, which are designated as “incorporated places”, had limited success as the populations of these regions are not directly accessible through the API.

### Sample set creation

The performance of ADCI-HT was evaluated by creating and processing replicate sample sets programmatically, comprised of fixed and variable numbers of distinct metaphase images and complete samples. While the scheduler is capable of handling samples of different sizes, sets of synthetic samples consisting of 500 images each were randomly selected from a larger pool of images exposed to the same radiation dose to estimate average processing speed per sample. This fulfills the minimum criteria for the dicentric assay [21]. To ensure that all images were processed, duplicate samples were also created by splitting consecutive subsets of 500 images into new samples. Residual subsets with fewer than 500 images were added to the last sample. Finally, samples of intermediate exposure levels were constructed by mixing sample pairs at different exposure levels from the same laboratory.

A large multi-laboratory exercise (40,000 samples) was simulated by duplicating 10 different exercise samples from five dosimetry laboratories in equal proportions (Canadian Nuclear Laboratories [CNL]: S02, S05, S09; Health Canada [HC]: S01, S05, S07, S08; Radiation Protection Center, Lithuania [LT]: 0.4Gy; Public Health England [PHE]: B; Dalat Nuclear Research Institute, Vietnam [VN]: 2.3Gy). These duplicates were dynamically split into sample sets based on the criterion that each set contains at most 25,000 metaphase images. Dose estimation for samples was performed using the ADCI-derived biodosimetry curve corresponding to the respective laboratory from which the samples were derived. For each run, the number of images processed in a sample set is related to the amount of available computer memory, as the output for processing all images is maintained in random access memory until the batch process is completed.

### ADCI-HT Resource allocation

In general, the CPU resources allocated were limited to 4 nodes by the default job queuing system. Priority scheduling was approved by system administrators to determine performance in a machine-optimized environment. Higher priority runs maximized the number of processors that could be simultaneously allocated in the supercomputer (though processes could still be delayed due to the BG/Q queuing system). These included 1 and 4 node runs of sample sets (100 runs with 400 samples each), with each node containing 1,084 cores (4 threads per core).

### Comparison with other systems

The performance of the high performance, ADCI-HT interpretation of sets of metaphase images was compared to fully automated (ADCI-Windows) and semi-automated analyses (DCScore [Metasystems]) on single computer systems. The DCScore estimates were based on Romm et al. 2013 [22], with a modification to include a missing processing step of performing thumbnail gallery review and selection of 500 optimal images. The time to perform this step was determined for 3 samples, averaged, and added to the time reported for the other review and processing steps.

## Results

This study attempts to determine the volume and capacity required to interpret biodosimetry tests by simulating provision of biodosimetry results for affected populations in high yield nuclear incidents. We used ADCI to carry out this task, which has been demonstrated to provide accurate dose estimates for samples of unknown exposure, based on IAEA compliant triage criteria. ADCI estimates radiation exposures using a fully automated process based on a calibration curve derived from the same biodosimetry laboratory [14–16]. The Windows-based Desktop version of ADCI was migrated to the PowerPC operating system of IBM BlueGene/Q (BGQ) as ADCI-HT, which significantly improved the throughput of these analyses.

Initially, the actual clock time (ACT) to perform large scale analyses of datasets consisting of samples, each consisting of 500 randomly selected metaphase images. To gain perspective, we processed 50 samples of 500 images each from CNL on both the Desktop and BGQ versions of ADCI. The processing time on Windows was 11h 10m, while on BGQ they required 30m 52s, which is equivalent to ADCI-HT being 21.7-fold faster than the Desktop version.

We also created and processed parallel sets of 120 samples, comprised of 15,970 images with ADCI-HT, by randomizing and splitting original calibration and exercise samples from 4 different laboratories. This involved submission of 54 computer runs of sample sets containing 120 samples, with each requesting 1 processing block (or 1,024 CPUs). With dedicated or high priority access to ADCI-HT and all runs ending successfully, the ACT would be equal to the maximum processing time of each run in parallel. In fact, ACT for all of these jobs ranged from 38m 37s to 1h 18m 2s, with an average of 1h 18s. As these jobs were submitted to the general queue (which is shared between all users of the system), the latency or wait for available CPU resources was variable, with the slowest set of samples to finish requiring 6h 6m 46s. The largest test involved 164 sets of samples, each set consisting of 100 samples (16,400 samples total), with each sample set submitted as a separate run. The maximum ACT to process one of these jobs was 1h 09m 58s, while the minimum was 30m 24s, averaging 45m 45s per run. The cumulative processing time without parallelization was 125h 2m 20s. We also compared these results with those from larger sized sample sets allocated with proportionally increased CPU resources. When requesting an allocation of 4 processing blocks (or 4,096 total CPUs; use of added resources required special permission), no discernable difference in ACTs was observed (maximum and minimum ACT per run were respectively 1h 5m 52s and 34m 9s, with an average time of 48m 48s). This was primarily due added queuing times when requesting more resources. Constraints on hardware architecture meant that allocation of proportionately increased CPU resources did not confer any advantage for processing larger sample sets.

### Benchmark of Population-Scale Nuclear Incident Simulations

HPAC-derived plumes were generated across 15 populated regions within the United States. The HPAC plume is represented as a series of topological contours (ovals) representing various levels of radiation (ranging from 1.0 Gy [the outermost ring] to 7.0 Gy [the innermost ring]). Figure 3 shows the radiation plume derived for the Boston scenario. Using U.S. sub-division boundary files and census information, we computed the overall population expected to overlap the >1Gy and >2Gy contours of the HPAC plume (Table 1; column 2 and 3, respectively). We then used sample data to simulate ADCI-HT analysis of all persons expected to obtain >2Gy exposure (500 images per patient). With 4 nodes with 16,384 total cores, the overall processing time ranged from 0.6 to 7.4 days (Burlington VT and Boston MA, respectively; Table 1) depending on population density.

**Figure 3.**
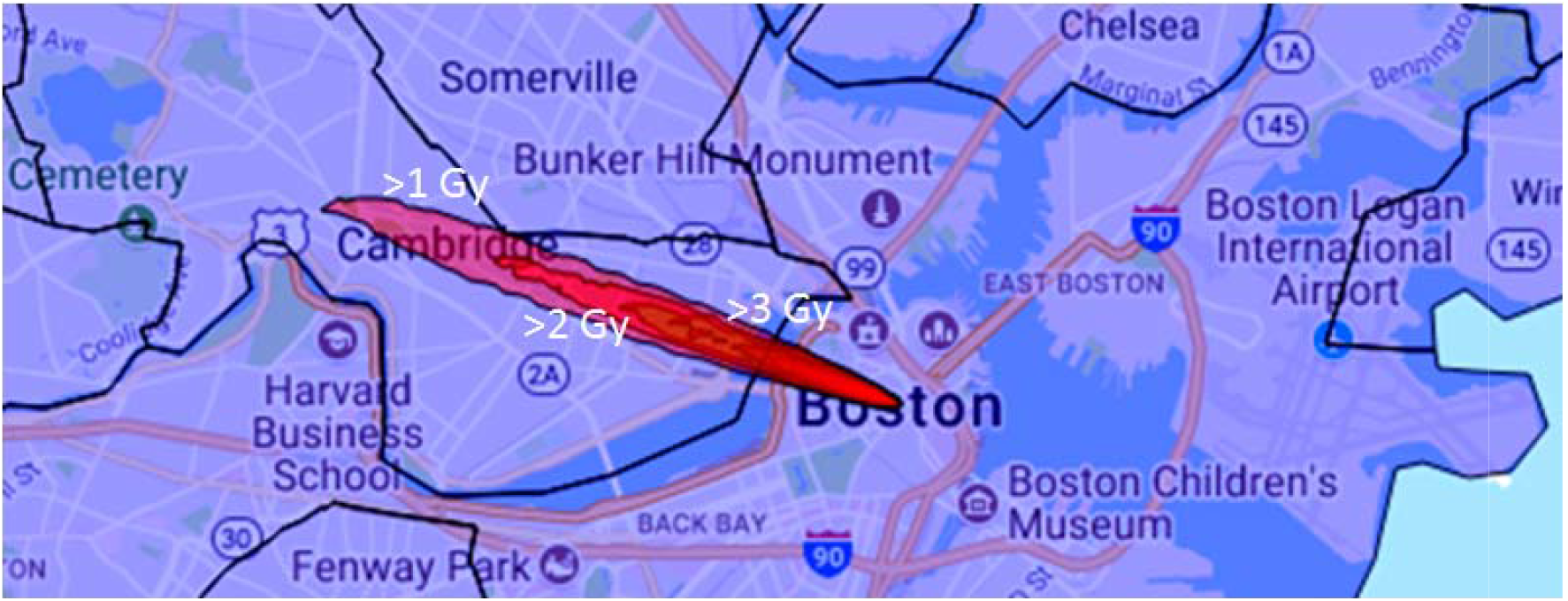
Capacity of ADCI-HT for dose estimation in a high yield mass casualty. HPAC-derived plume presented intersects with contours of >2 Gy include Boston, Suffolk County, Massachusetts (population = 673,184) and Cambridge, Middlesex County, Massachusetts (population = 110,651), leading to a total population of 783,835. Extrapolating from 6,480 sample run (1.3 hr for 4 nodes), the time required for ADCI-HT to analyze samples from the total population would be 157.25 hours or 6.5 days.

**Table 1.** Estimated exposed populations in US metropolitan and BG/Q processing times after high yield nuclear incident^.

| City State | No. persons with<br>> 1 Gy exposure | No. persons with<br>>2 Gy exposure | Processing time*<br>(>2 Gy) in days |
| --- | --- | --- | --- |
| Burlington VT | 92,008 | 61,231 | 0.58 |
| Portland ME | 102,720 | 66,937 | 0.63 |
| Manchester NH | 136,356 | 110,506 | 1.04 |
| Boston MA | 865,157 | 783,835 | 7.37 |
| Albany NY | 98,111 | 98,111 | 0.92 |
| Buffalo NY | 256,902 | 256,902 | 2.41 |
| Hartford CT | 309,626 | 216,569 | 2.04 |
| New Haven CT | 138,776 | 129,934 | 1.22 |
| Stamford CT | 191,472 | 191,472 | 1.80 |
| Newark NJ | 476,853 | 281,764 | 2.65 |
| Camden NJ | 230,942 | 110,043 | 1.03 |
| Harrisburg PA | 78,870 | 73,817 | 0.69 |
| Pittsburgh PA | 319,221 | 303,625 | 2.85 |
| Detroit MI | 672,795 | 672,795 | 6.32 |
| Grand Rapids MI | 350,236 | 196,445 | 1.85 |
| Milwaukee WI | 595,047 | 595,047 | 5.59 |

\*based on 16,384 cores, 4 nodes (non-prioritized jobs scheduled); <sup>^</sup> 5 megaton airborne yield, plume based on historically prevalent wind speed and direction

### Performance of ADCI-HT on Sample Sets from Multiple Laboratories

The volume of samples in a large-scale nuclear incident would likely exceed the capacity of any individual biodosimetry laboratory to process. We simulated processing and dose estimation of 40,000 samples (1,892 sample sets) by 5 laboratories, each of which has a distinct dose calibration curve. ADCI-HT took 25h 8m 5s to finish processing the simulated samples, i.e. each duplicated from the same set of metaphase images. These cumulative set of samples contained 46,196,000 metaphase cell images in total and were processed with 37,888 CPUs on average (Figure 4). The respective durations for processing of the sample sets from the different laboratories (CNL: 14h 6m 22s; HC: 6h 27m 22s; LT: 3h 16m 21s; PHE: 1h 28m 40s; VN: 1h 44m 2s) were proportionate to the total image counts in each set of samples. The dose estimates for individual samples produced by the ADCI-HT version were identical to those produced by the Desktop version on all these samples (not shown), except those from HC. The deviation are due to differences between the order of images selected by the ‘Group Bin Distance’ filter used in the HC dataset (which selected the top 250 images; [14]), where multiple images had the same value, resulting in the insignificant discrepancies in the dose estimates generated by the Desktop and HT versions of ADCI.

**Figure 4.**
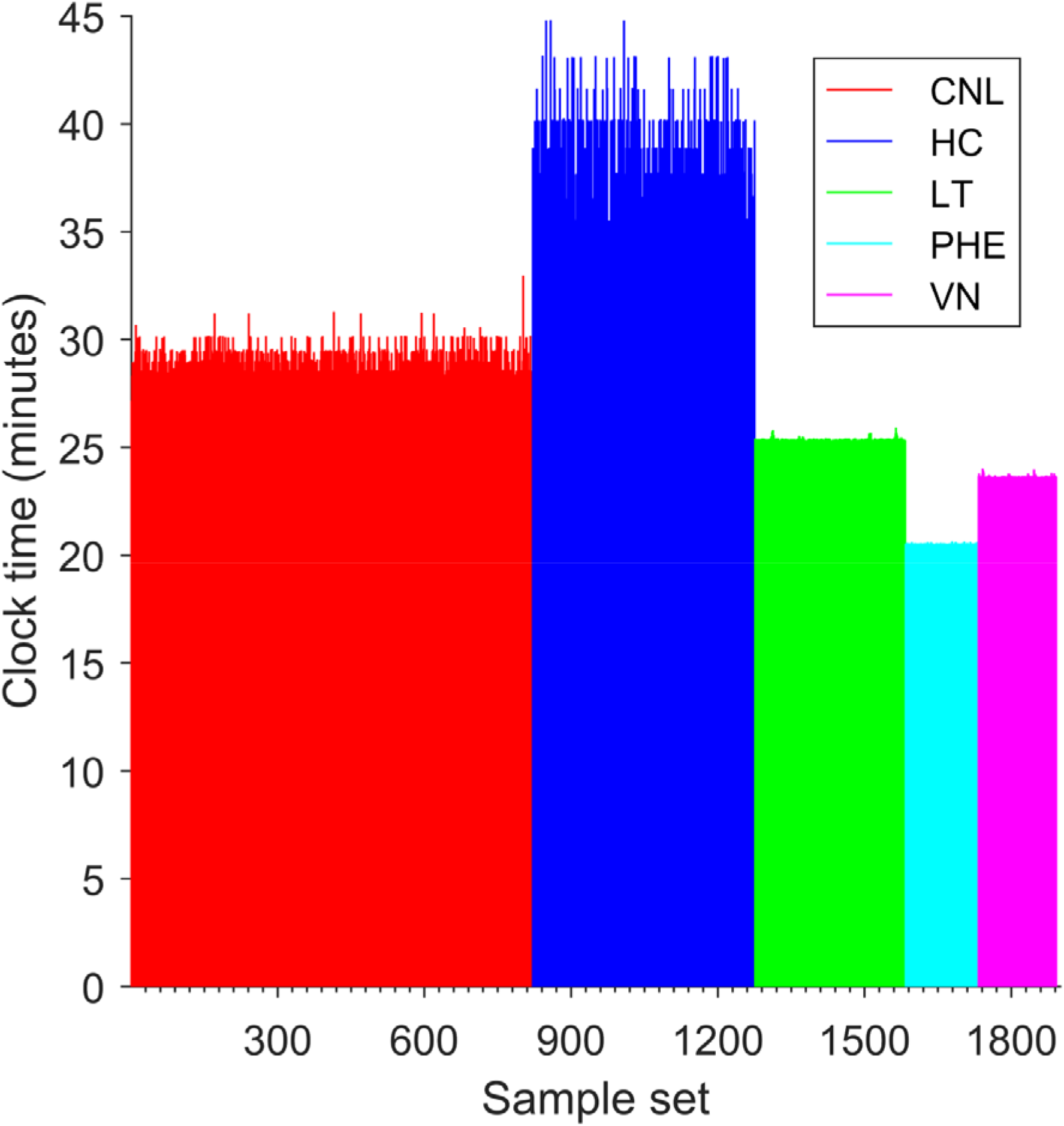
Large scale simulation based on multi-centre biodosimetry exercise sample sets. Test image samples acquired from 5 separate laboratories consisting of 46,196,000 metaphase images were processed by ADCI-HT in just over 25 hours. Duration of each sample set are as follows: CNL: 14:06:22; HC: 06:27:22; LT: 03:16:21; PHE: 01:28:40; VN: 01:44:02. Times correlated to the total number of images in each data set.

### Comparison of ADCI-HT with ADCI-Windows and DCScore performance

Dose estimation was much less computationally expensive with ADCI than with a semi-automated software product, DCScore (Metasystems), which requires manual curation of images and confirmation of predicted DCs. To determine whether ADCI-HT could provide timely dose estimates with available cytogenetic data for a moderate sized population of potentially exposed individuals, we compared the processing requirements for samples by ADCI-Windows and HT versions with DCScore. Results for individual samples were extrapolated to a population consisting of 1,000 samples (Table 2). The differences in time to process a single sample were negligible between these platforms. Only the HT version was able to estimate exposures with sufficient speed for the population-based analysis to triage all patients to effectively treat them. However, multiple instances of ADCI-Windows running on separate laptop computers (n=2) would be required to fulfill assessment of this population in 1 day (Figure 5). DCScore is unsuitable for this task, as it would require nearly a month to analyze a population of this size, due to manual review and selection of metaphase cells in each sample and confirmation of candidate dicentric chromosomes.

**Table 2.**
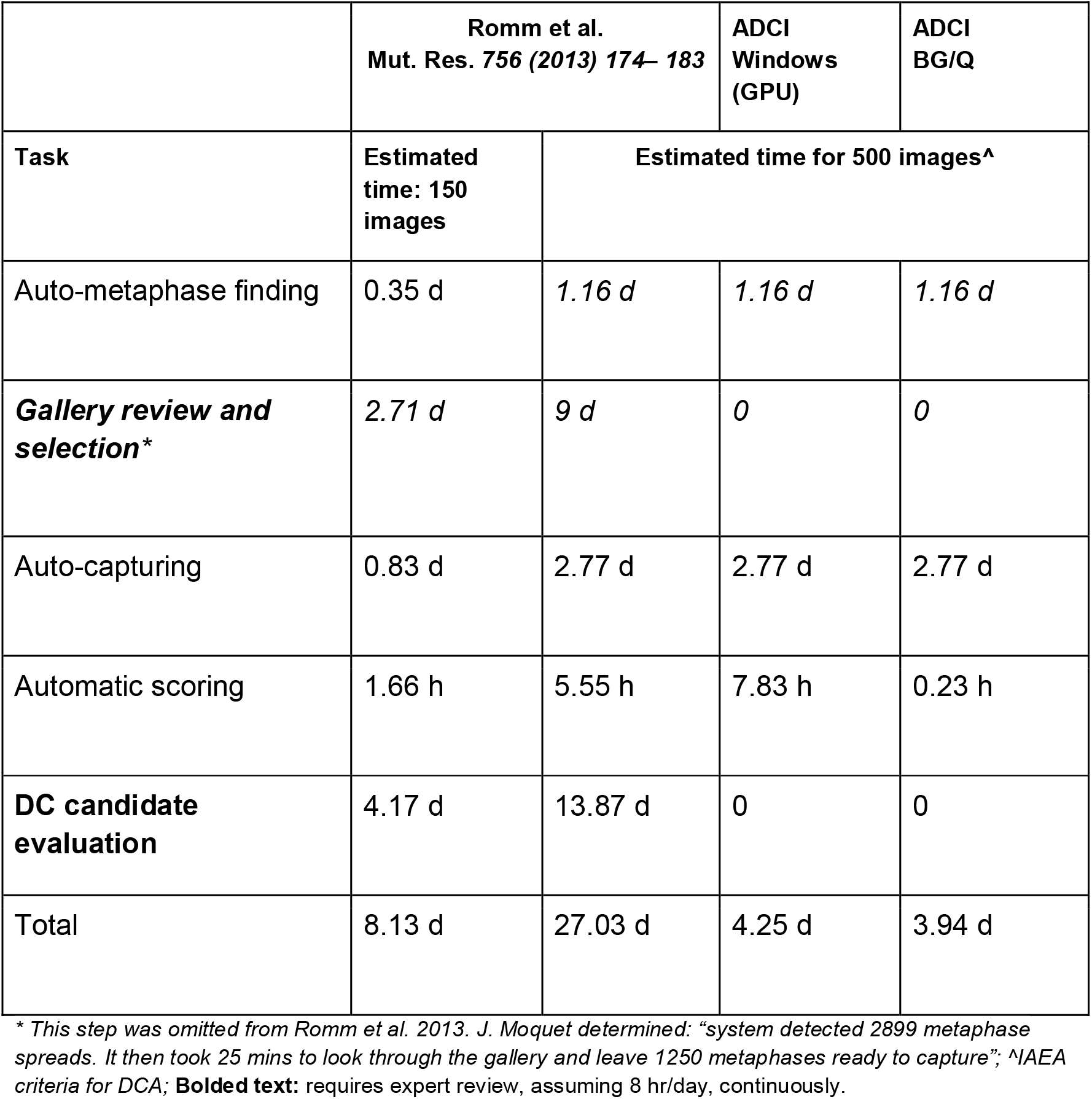
Comparison of capture and determination of DC counts in 1000 samples. Assumes 10 microscope capture systems for capture and either 10 DCScore or 10 ADCI–Desktop or 1 ADCI-BGQ) systems

**Figure 5.**
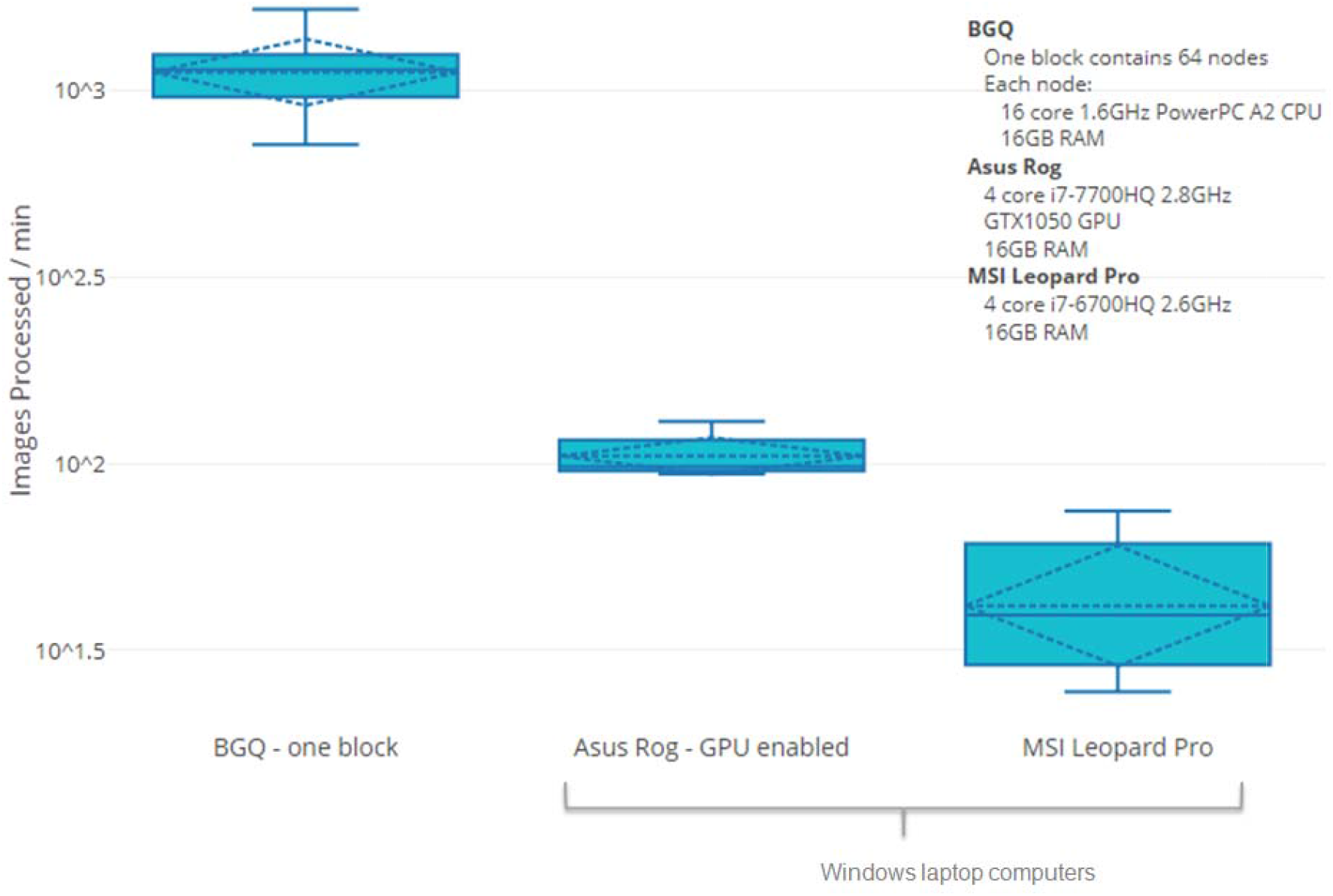
Comparison of ADCI throughput for different hardware platforms. The rate in which images are processed using only a single BG/Q processor block is ~10x fold faster than a GPU-enabled i7 laptop (4-core i7-7700HQ 2.8GHz processor, GTX1050 GPU and 16GB RAM). Image processing throughput can be increased further using multiple processor blocks via BG/Q scheduler.

## Discussion

ADCI-HT has the capacity to analyze large quantities of cytogenetic biodosimetry data which may be generated after a large radiation event. Laboratory capacity to process samples and generate metaphase image data has benefitted from automation. With a distributed testing model, it would be feasible to generate different sets of metaphase image samples in multiple laboratories, each customized to use their own image selection models. The samples would be uploaded and images processed in parallel via a cloud-based system running ADCI-HT, which would estimate exposures for all samples either simultaneously or possibly in large batches in rapid sequence. We suggest that leveraging existing laboratory infrastructure and supplanting limitations in any of these elements with existing automation at large commercial cytogenetic laboratories may be able to fulfill future testing demands for biodosimetry testing in a population scale nuclear event or accident.

Rapid triaging using fewer metaphase images or other assays (micronuclei, biomarkers in urine, miRNA, or gene expression) have been proposed as alternatives to the DCA, however these approaches may not be sufficiently accurate to robustly distinguish individuals eligible for treatment (>2 Gy exposures; [23]). Rapid bioassays have been developed for triage evaluation of potentially exposed populations, however specific frameworks have not been demonstrated by large scale implementation of these tests. Throughput estimates have largely been theoretical and based on the performance of the assays themselves, extrapolated to large populations. The details regarding equipment capacity, availability of sufficient quantities of critical reagents and trained personnel to perform and interpret these assays have not been addressed in the Concept of Operations [24].

HPAC provides the distribution of the physical dose in a certain region. However, the physical doses may not be identical to the biological doses but may, in some instances, be correlated [25]. Spectral clustering [26–28] and geostatistics [29] can be used to determine the minimum number of data points (e.g. locations in the map) that are needed to construct similar distribution. However, the complexity of the calculations needed for spectral clustering become increasingly prohibitive with large datasets. Our efforts have turned to geostatistical estimation of radiation exposures.

ADCI-HT could be an instrumental resource in the rapid identification of patients requiring treatment in population-scale nuclear incidents. Depending on population-density, ADCI-HT software can identify DCs and estimate dose of a population within an irradiated zone in 0.6-7.4 days (Table 1; Figure 3). ADCI-HT by itself, however, only accelerates image processing and DC identification. The acquisition of said samples, the preparation of metaphase cells and the capture of DC images makes the testing of entire populations logistically impossible. Sampling and testing requirements can be reduced via geostatistical sampling; a method of estimating the spatial boundaries of a region using a small subset of samples at various locations. Applying such methods can limit the number of patients requiring testing while reducing sampling time required for first responders, expediting identification of those requiring treatment in scenarios where time and resources are limited.

Despite this significant reduction in processing time using ADCI software, there are still potential improvements that may increase the feasibility of implementing this software as a routine biodosimetry laboratory resource. Although ADCI-HT was benchmarked as faster than Desktop ADCI, access to supercomputers may not always be feasible, and therefore increasing the rate of analysis via conventional ADCI is still crucial. We have recently implemented GPU acceleration on the Desktop platform. Preliminary results showed a ~8x increase in sample processing speed, which may not be as fast as ADCI-HT but may be adequate in some population-scale scenarios. We are also implementing the contaminated Poisson method to calculate the mean DC frequency within the irradiated fraction of partially irradiated samples.

We have presented a high-throughput implementation of ADCI which rapidly processes metaphase images and provides radiation dose estimations. The dose estimates results were concordant with those generated by the Desktop version when analyzing exercise sample data from 5 different laboratories. This level of dedicated computing resources available for the 40,000 sample run made a significant difference in obtaining timely dose estimates for the entire population, as a single processing block would have been insufficient (i.e. 37 days of ACT) for timely estimation of population-scale exposures. ADCI-HT therefore achieves adequate performance to deliver timely and accurate dose estimates in a mass casualty radiation event.

## Acknowledgements

We are grateful to the SOSCIP Consortium (J.H.M.K., P.K.R.), Natural Sciences and Engineering Research Council of Canada (Engage Program; E.W.), Ontario Centres of Excellence (Talent Edge Postdoctoral Fellowship Program; J.H.M.K., P.K.R.), CytoGnomix (P.K.R.) for support of this project. Certain data used in this study were obtained as part of Coordinated Research Project E35010: Applications of Biological Dosimetry Methods in Radiation Oncology, Nuclear Medicine, and Diagnostic and Interventional Radiology (MEDBIODOSE), which was carried out under the sponsorship of the International Atomic Energy Agency.

## Supplementary Figure

**Supplementary Figure 1.**
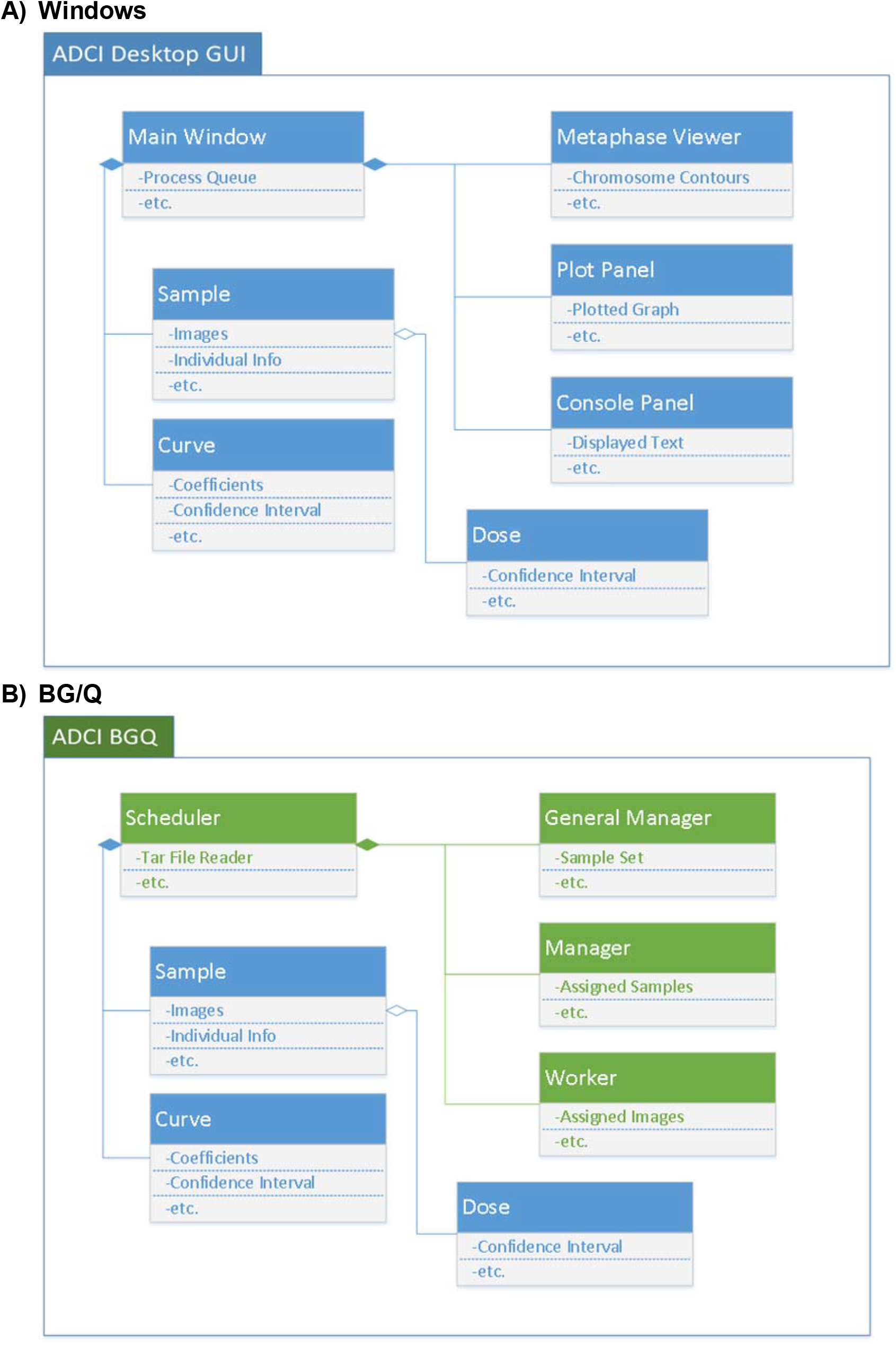
Adapting ADCI Windows Desktop Graphical User Interface to ADCI-HT on BG/Q. Data flow diagram illustrating the steps required to perform metaphase image processing tasks using A) Windows ADCI, and B) BG/Q ADCI (ADCI-HT) software platforms.

